# Song preferences predict the quality of vocal learning in zebra finches

**DOI:** 10.1101/2021.06.01.446570

**Authors:** Carlos Antonio Rodríguez-Saltos, Aditya Bhise, Prasanna Karur, Ramsha Nabihah Khan, Sumin Lee, Gordon Ramsay, Donna L. Maney

**Affiliations:** Department of Psychology, Emory University, Atlanta, GA 30322 USA; Marcus Autism Center, Children’s Healthcare of Atlanta, Atlanta, GA 30329 USA; Department of Pediatrics, Emory University School of Medicine, Atlanta, GA 30329 USA

**Keywords:** birdsong, social learning, social reward, songbird

## Abstract

In songbirds, learning to sing is a highly social process that likely involves social reward. Here, we hypothesized that the degree to which a juvenile songbird learns a song depends on the degree to which it finds that song rewarding to hear during vocal development. We tested this hypothesis by measuring song preferences in young birds during song learning and then analyzing their adult songs. Song preferences were measured in an operant key-pressing assay. Juvenile male zebra finches (*Taeniopygia guttata*) had access to two keys, each of which was associated with a higher likelihood of playing the song of their father or that of another familiar adult (“neighbor”). To minimize the effects of exposure on learning, we implemented a reinforcement schedule that allowed us to detect preferences while balancing exposure to each song. On average, the juveniles significantly preferred the father’s song early during song learning, before they were themselves singing. At around post-hatch day 60, their preference shifted to the neighbor’s song. At the end of the song learning period, we recorded the juveniles’ songs and compared them to the father’s and the neighbor’s song. All of the birds copied father’s song. The accuracy with which the father’s song was imitated was positively correlated with the peak strength of the preference for the father’s song during the sensitive period. Our results show that preference for a social stimulus, in this case a vocalization, predicted social learning during development.

## INTRODUCTION

Conspecific signals are attractive to receivers. Research on this attraction has focused mostly on signals such as courtship displays (Andersson, 1994) or foraging calls (Suzuki & Kutsukake, 2017). These signals share in common that attending to them may provide immediate benefits to the receiver. However, being attracted to conspecific signals is also important when the benefits are not immediate. Consider vocal learners, such as seals, bats, and songbirds (Janik & Slater, 1997; Jarvis, 2006). Juveniles of these species must attend to signals, such as speech or song, in order to imitate them; the benefits of doing so are often seen in adulthood, once these signals can be used to share information (Suzuki & Kutsukake, 2017), attract a mate, or secure a territory (Catchpole & Slater, 2008). Thus, learning these signals likely depends on being attracted to them at a time when those benefits of signaling are beyond reach (Rodríguez-Saltos, 2017). There should, therefore, be a great deal of selection pressure on young learners to be attracted to these signals, even in the absence of immediate mating opportunities or food rewards.

Songbirds lend themselves well to studying the processes by which attraction to song contributes to vocal learning. Zebra finches (*Taeniopygia guttata*), which are among the most commonly studied songbirds in the lab, are actively engaged in learning to sing. They memorize songs during social interactions with adults, and their degree of attention towards adult song “tutors” during these interactions predicts the quality of song imitation (Chen et al., 2016). Juvenile finches are easily lured to press keys that elicit playback of song, and if given the opportunity, they will elicit playback hundreds of times per day. The fact that young finches are willing to work to elicit playback of song shows that this stimulus is rewarding to them, just as access to food or mates is rewarding to animals that are willing to press levers to obtain them. We hypothesize that the degree to which a particular song is rewarding dictates the degree to which it is learned.

Song learning is thought to rely on a mechanism similar to imprinting (Baran, 2017) in that it occurs during an early critical period and requires social experience. In zebra finches, only the males sing. Beginning at around 20 days post-hatch (dph), before they are able to sing, young pupils select an adult male tutor and begin to memorize his song. The chosen tutor is nearly always the father. Once a juvenile has early experience with the father, if the father is removed before song memorization is complete, the juvenile will choose a tutor that looks like the father (Mann et al., 1991) or sounds like him (Clayton, 1987). If a juvenile is reared from hatch by a foster male, he will learn the song of that male even if the biological father’s song is also heard in the room (Eales, 1987; Immelmann, 1969). The need for social interaction during learning is further evidenced by the fact that young finches do not imitate songs that are presented via passive playback; finches that have been passively tutored produce songs that are similar to those of finches that have not been exposed to song at all (Adret, 1993; Derégnaucourt et al., 2013). Together, these studies show that the learning process is strongly shaped by social contexts that may make one potential tutor’s song more rewarding to hear than another’s. In other words, early experience with the father is likely to lead to preferences for the father’s song, which could lead to enhanced learning of that song.

Preference for the song of the father, in the absence of the father himself, has been studied in finches both by measuring phonotactic responses and by using operant conditioning techniques. In the Bengalese finch (*Lonchura striata domestica*), a species that is closely related to the zebra finch, juveniles more frequently approached speakers playing the father’s song than speakers playing unfamiliar songs (Fujii et al., 2021). This preference may result from the father’s song being more rewarding than other conspecific songs. It is still unclear, however, whether preference for rewarding songs contributes to the learning of those songs. Adret (1993) used a key-pressing assay to detect preferences in adult male zebra finches that had already finished learning to sing (Adret, 1993). These adults preferred the songs with which they were tutored over the songs of other adults. Although this result is intriguing, it does not indicate whether preferences in adulthood reflect preferences during learning. Importantly, we do not know whether such preferences predict learning.

It may seem logical that birds learn their favorite songs best. This prediction is difficult to test, however, because in traditional preference assays, preference is confounded with exposure. If an animal chooses one stimulus over another, indicating preference, that animal exposes itself to the preferred stimulus more than the non-preferred one. Because exposure can affect learning (Tchernichovski et al., 1999), we developed a novel reinforcement schedule that allowed us to measure preference while balancing exposure. Using a key-pressing assay in juvenile male zebra finches, we tested preferences for the father’s song over that of a familiar, unrelated male. We measured preference each day beginning at nutritional independence, before the juveniles began singing, and ending at song crystallization, when the song is fully learned and does not change thereafter (Fig. 1A; Johnson et al., 2002; Johnston, 1988; Kollmorgen et al., 2020; Tchernichovski, 2011). At that point, we recorded the adult song to test our hypothesis that early preferences for the father’s song would predict learning of that song.

**Fig. 1.**
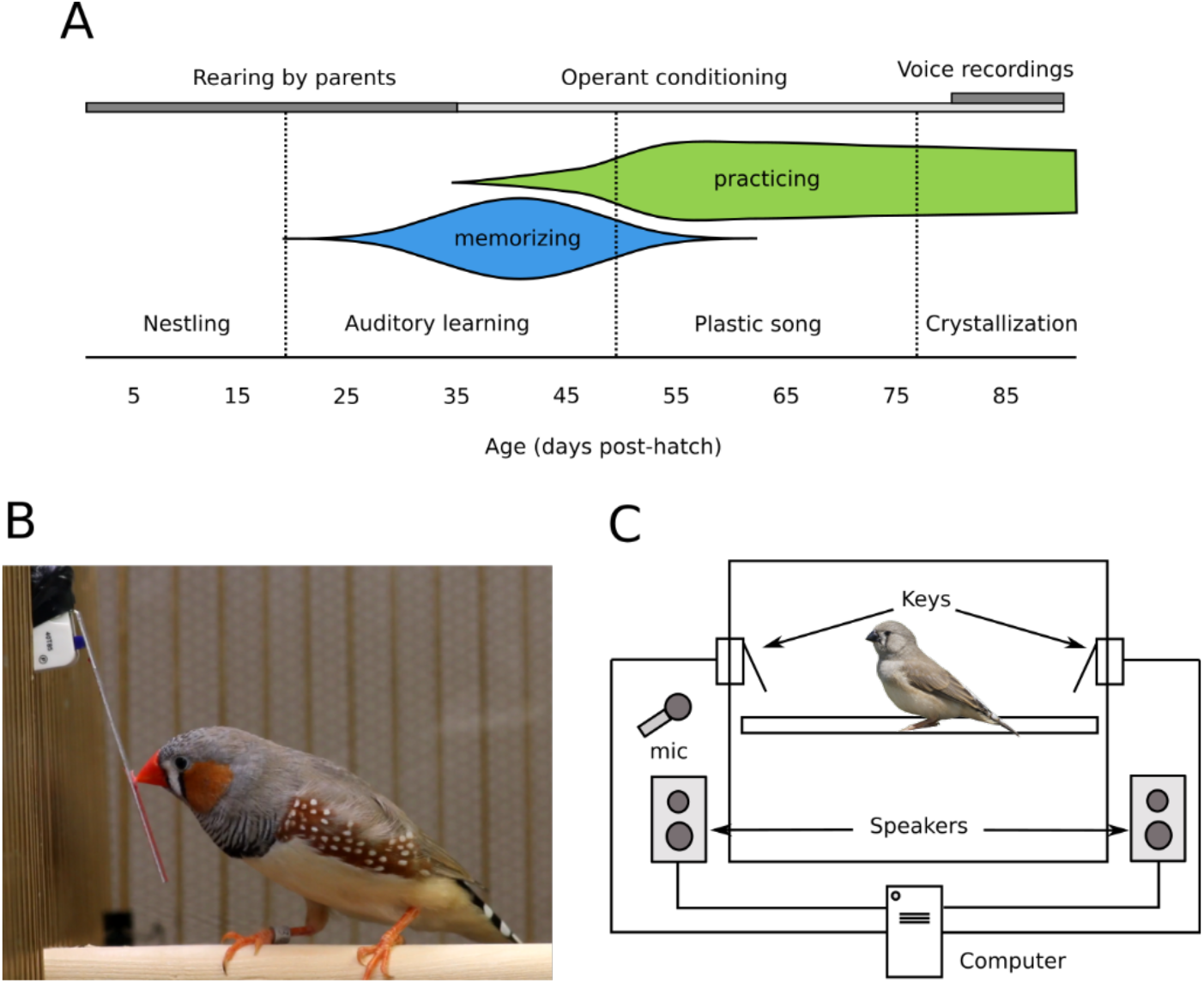
Experimental set-up. (A) Experimental timeline (top) and developmental trajectory of song learning in male zebra finches. Zebra finches were reared by their parents in a room where they could also listen to, but not see, adult male zebra finches other than the father (“neighbors”). At approximately 35 days post-hatch (dph), when finches are nutritionally independent (Zann, 1996), the juveniles were transferred to an operant chamber equipped with keys that were associated with playback of father’s or neighbor’s song. Preference for father’s or neighbor’s song was measured daily while the bird remained in the operant chamber, until 90 dph. This timeline of operant conditioning covers most of the developmental trajectory of song learning. At 35 dph, finches are in an auditory phase of learning, during which they memorize adult song (Eales, 1985) and start practicing singing (Zann, 1996). Singing rates start increasing by age 50 dph. Approximately at the same time, they enter the “plastic song” phase (Johnson et al., 2002; Kollmorgen et al., 2020), during which some memorization may still occur but ends by 65 dph (Eales, 1985). Finally, song crystallization begins by 77 dph and finishes by 90 dph (Johnson et al., 2002; Zann, 1996). In the crystallization phase, between 80-90 dph, we recorded vocalizations of the juveniles and compared them to father’s and to neighbor’s song. (B) An adult zebra finch presses a key in an operant chamber to elicit playback of conspecific song. The same setup was used with the juveniles in our experiment. Photo by CAR-S. (C) The operant chamber consisted of a 14×15×17 inch cage, inside which two keys were placed on opposite walls. One key was associated with playback of father’s song and the other with playback of neighbor’s song. Outside the cage, one speaker assigned to each key played the songs. Photo of finch by Lip Kee Yap, shared under the Creative Commons Attribution-Share Alike 2.0 Generic license.

## RESULTS

### Developmental trajectories of song preference

Juvenile male zebra finches, or “pupils”, were reared by their parents, with full access to the father, until nutritional independence (median age 37 dph). They were then transferred to operant cages containing two keys (Fig. 1B, 1C). One of the keys was more likely to play the father’s song (hereafter referred to as “father’s song”). The other key was more likely to play the song of a different male (hereafter referred to as “neighbor’s song) that had been housed in the same room, but not the same cage, as the juvenile’s family cage and which had a song rate similar to the father. Each day, the pupil could press the keys to trigger 30 playbacks of each song. Because of our novel reinforcement schedule (Fig. 2; see Methods), exposure to each song was balanced each day. Each pupil heard 30 playbacks of father’s song and 30 of neighbor’s song, then the keys would trigger no further playbacks until lights-on the following day.

**Fig. 2.**
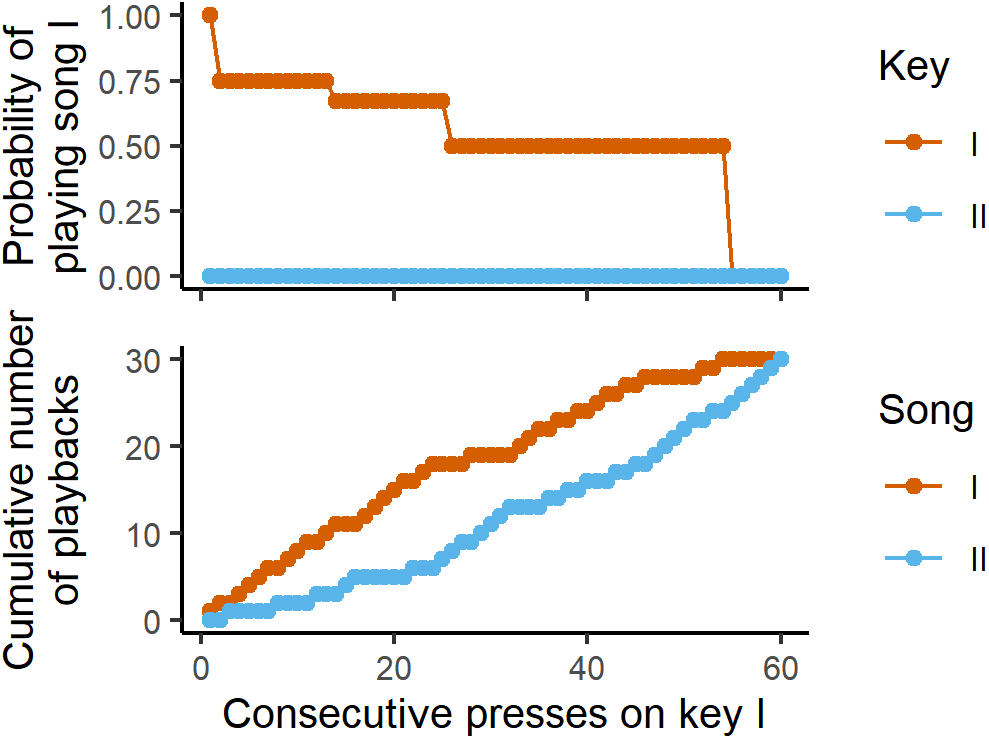
Reinforcement schedule to detect song preference while balancing exposure. A bird is presented with keys I and II, which are associated with a higher probability of playing song I and song II, respectively. Here we present an extreme example to illustrate how the schedule is able to balance exposure despite a strong preference. In this scenario, the bird prefers song I and presses key I only. The probability of playing song I by pressing that key is high at the beginning of the session (A), to help the bird form the association between that key and the song. As the bird keeps pressing key I, the probability decreases step-wise from 1 to 0.5, to prevent song II from lagging far behind in the playback count (B). This decrease, however, is not enough to balance exposure, and therefore, if the bird switches keys, key II plays only song II until the playback count of song II is balanced with song I. After enough presses on key I, song I eventually reaches a quota of 30 playbacks and ceases to be played. Afterwards, only song II is played, until that song also reaches the quota. Importantly, there is always a large difference between the keys with respect to the probability of hearing the preferred song. When the key associated with preferred song is playing that song only 50% of the time, the other key plays non-preferred song 100% of the time.

After being transferred to the operant cages the pupils engaged with the task and were exhausting the daily quota of 60 playbacks (30 of each song) within 5.0 ± 3.72 days (mean ± standard deviation). The age at which they started exhausting the quota did not correlate with the age at which they were transferred to the operant conditioning cages (Spearman’s rho = −0.32; *p* = 0.31). Pupils remained with the operant conditioning task until a median age of 89.50 dph (IQR: 80.90-90.00). One juvenile, which did not exhaust the quota of both songs after 13 days of being housed in the cage, was excluded from the study, leaving n = 12 that completed the keypressing portion. Their crystallized songs were recorded between 80-90 dph for comparison with the song playbacks.

On the basis of previous research (Clayton, 1988; Fujii et al., 2021; Miller, 1979; Riebel et al., 2002) we hypothesized that the juveniles would show a preference for father’s song over neighbor’s song, and we therefore calculated preference with respect to father’s song (preference for neighbor’s song is simply the preference for father’s song subtracted from 1). For the 12 pupils that reliably exhausted the quota each day, we estimated the strength of the preference for father’s song over neighbor’s song daily by calculating the proportion of presses on the key associated with father’s song. Scores above 0.5 indicate that father’s song was preferred over neighbor’s song. On average, the father’s song was preferred over neighbor’s song between ages 45-52 dph (preference for father’s song > 0.5; 0.5 outside 95% confidence interval [CI]) (Figure 3A), during the auditory phase of learning. The neighbor’s song was preferred between developmental ages 57-65 dph and then again between 87-90 dph (preference for father’s song < 0.5; 0.5 outside CI). The shift from preferring father’s song to then preferring neighbor’s song was also seen in many individual developmental trajectories (Fig. S2).

**Fig. 3.**
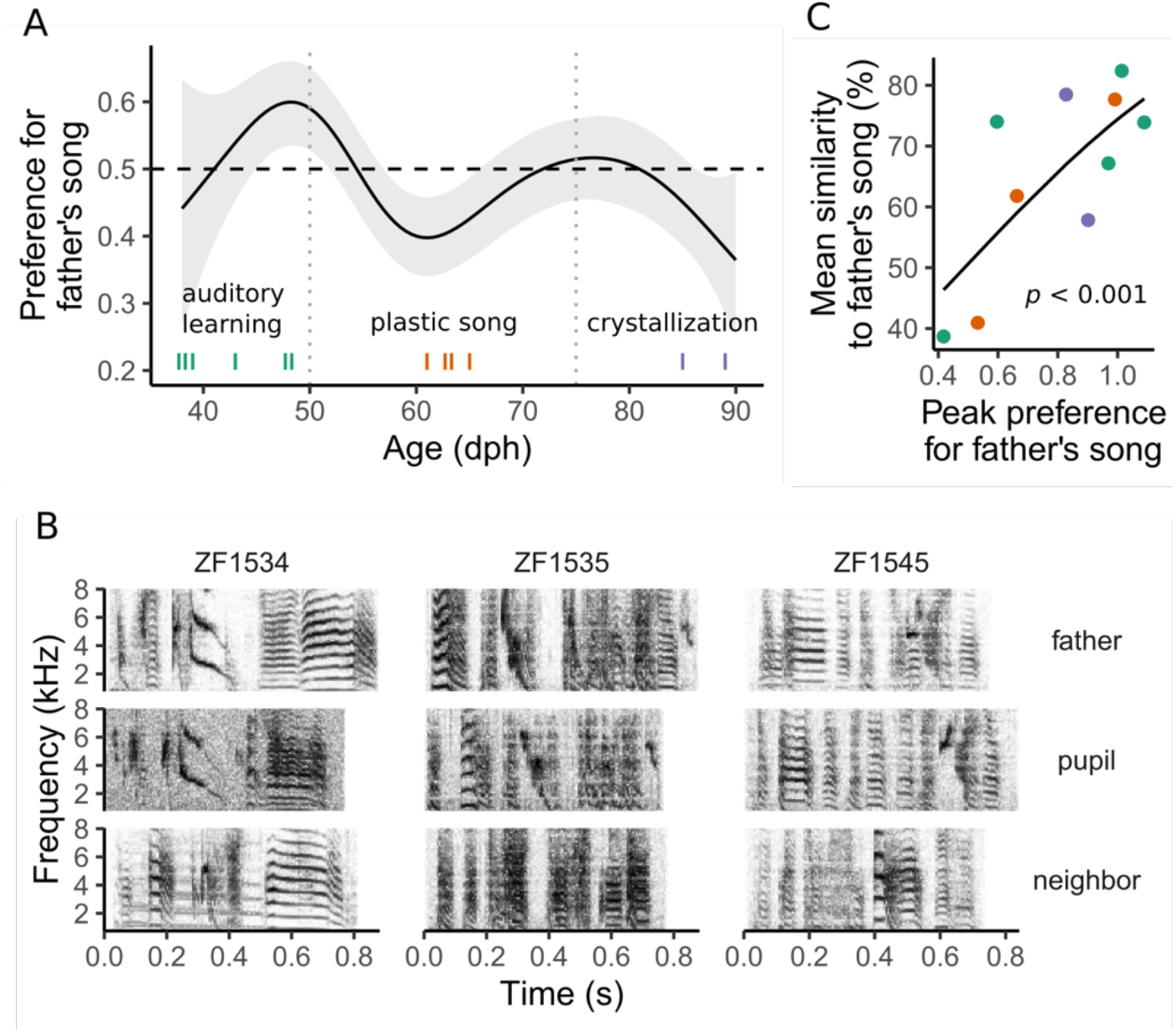
Relationship between the preference for father’s song and imitation of that song. A) The plot shows the average trajectory of the preference for father’s song. The shaded area indicates the 95% confidence interval. The horizontal dashed line indicates no preference for either song. Vertical dashed lines indicate boundaries between developmental phases (see Fig. 1A). Ticks near the bottom of the plot indicate the age at which individual birds reached their peak preference for father’s song (for individual trajectories, see Fig S2). On average, preference peaked during the auditory learning phase. B) Peak preference for father’s song was significantly correlated with mean similarity to that song. The curve is the fit of a beta regression. Ticks in (A) and dots in (B) are colored according to phase in which the bird reached maximum preference for father’s song. C) Spectrograms of the songs of three pupil exemplars, shown next to the spectrograms of the songs of their respective father and neighbor. Syllable shapes in the spectrograms indicate that these three pupils imitated father song and not neighbor song. All other pupils also imitated father song or, in the case of one individual, did so most of the time (see text). Spectrograms were generated in R (R Core Team, 2020) using Hanning windows containing 512 samples each (11.61 ms for recordings sampled at 44100 kHz), with 50% overlap between windows.

For each pupil, we identified the maximum value on a smooth trajectory of preference for father’s song. For half of the pupils (6 out of 12), the maximum preference for father’s song was above 0.91. Except for one, all pupils had a maximum preference for father’s song above 0.5, which means that at that point in their trajectory, they preferred father’s song over neighbor’s song. For half of the pupils (6 out 12), the point of maximum preference for father’s song occurred before 50 dph, in the auditory phase of learning (Fig. 3A). For four pupils, it occurred in the plastic song phase and for only two birds, during crystallization (Fig. 3A). Maximum preference for father’s song was not correlated with the age at which it was reached (Spearman rho: −0.30, *p* = 0.35).

### Correlation between preference and imitation

To test whether the strength of the preference for a song predicted the quality of imitation of that song, we next quantified imitation by examining the similarity between each pupil’s song and the song it chose to copy. On the basis of visual inspection of shared syllables, the songs of all pupils matched the father’s song (Fig. 3C, S3). The song of just one pupil also contained a single syllable from neighbor’s song (Fig. S4). Thus, we are confident that all of the juveniles in our study imitated father’s song exclusively or nearly so.

We next quantified the degree of similarity of pupils’ song to the father’s song using SAP2011 (Tchernichovski et al., 2000; Tchernichovski, 2011). The mean similarity score, per bird, was 65.3 ± 1.5 (mean ± s.d.). To test whether the strength of the preference for the father’s song predicted the quality of imitation of that song, we fitted beta regression models to our data on maximum preference and average similarity, controlling for brood as a random effect. Because technical issues prevented us from recording the crystallized songs of two pupils, our final sample size for this analysis was n = 10. We found a large, significant correlation between maximum preference for father’s song and average similarity of pupil’s song to father’s song (R^2^ = 0.51; *p* = 0.002, Fig. 3B).

### Relative importance of live tutoring vs. operant tutoring

Two birds not included in this study were tested during operant conditioning using two neighbor songs. We compared their crystallized songs to that of their father and the two neighbors. Despite not having had exposure to their father’s song after 35dph, and hearing only neighbor’s song during operant conditioning, the songs of these juveniles were similar to father’s song (Fig. 3C, S3). We did not find syllables from the two neighbor’s songs in the pupils’ songs.

## DISCUSSION

### Preferences and vocal learning

In this study, we showed that juvenile male zebra finches, when socially reared with the father until 35 dph, prefer to hear father’s song during the auditory phase of song learning and ultimately sing father’s song as well. Our finding that the pupils preferred father’s song is consistent with other studies using a variety of methods of assaying preference (Adret, 1993; Riebel et al., 2002), and in particular with a recent study showing that the preference for tutor song may peak early during vocal development (Fujii et al., 2021). Our result that all of the pupils imitated the father is consistent with other reports suggesting that early social experiences are critically important for tutor choice and high-quality song learning (Clayton, 1987; Mann et al., 1991; Mann & Slater, 1995; Williams, 1990; Zann, 1996).

Our most important finding is that the peak preference for father’s song strongly predicted learning of that song (Fig. 3B). Our interpretation of this finding is that the incentive salience of a song stimulus, in other words the pupil’s willingness to perform work to hear it, may facilitate vocal learning. In this study, the development of song preference was pupil-driven, not driven by exposure. Because exposure to father’s versus neighbor’s song was balanced in our paradigm, we conclude that the decisions about what was most attractive and what to learn were made by the pupil. We do not know whether a preference for father’s song represented an attachment to the father himself or simply a desire to hear the song that the pupil wished to learn. In fact, because positive feedback from the mother to the father may be perceived by the pupil (Carouso-Peck & Goldstein, 2019; Mann & Slater, 1994), we cannot say whether the father himself played any role in the development of preference for his song. Regardless of the mechanism, however, it is clear from our results that the strength of the juvenile’s preference predicted the quality of learning.

Vocal learning in songbirds has been hypothesized to be facilitated by the naturally rewarding properties of conspecific song, particularly the song of a caregiver during early development (Baran, 2017; Maney, 2013; Maney & Rodriguez-Saltos, 2016). Chen et al. (2016) recently showed that in zebra finches, attention to the tutor, defined as quietness and stillness at times when the tutor is singing, predicted the quality of learning. Our current findings are consistent with this idea and suggest further that this attentiveness may be related to the pupil’s attraction to the tutor’s song. These results may have interesting implications for language development in humans. Like song learning in songbirds, language learning in humans depends critically on early social interactions. It has been hypothesized that the motivation to attend to social stimuli plays an important role in learning speech. For example, gaze following, the desire to imitate a caregiver, and joint attention between a child and a caregiver predict language acquisition (reviewed by Kuhl, 2007). Alterations in social orienting, which likely indicate alterations in social reward, are thought to cause children with autism to develop language more slowly relative to typically developing children (Chevallier et al., 2012; Mundy & Burnette, 2005). Thus, zebra finches may serve as important models for the role of social reward in language learning.

### Developmental trajectory of song preferences

In this study, pupils showed a highly significant preference for father’s song during the auditory phase of song learning, that is, before ∼50 dph. During this phase, pupils begin to perform subsong, an early form of singing characterized by high variability and lack of distinguishable phrases (Zann, 1996). The critical process going on during this phase is likely not vocalizing, but listening; previous work in this species has shown that pupils engage in the most effective song memorization at around 45 dph (Deshpande et al., 2014), a time that, in our study, corresponded with a clear peak in preference for father’s song (Fig. 3A). Thus, the peak in memorization of a song happens at around the same time as a peak in the incentive salience of that song.

Beginning around 50 dph, the birds in our study showed a dramatic change in their song preference. On average, at that time they began to show a preference for neighbor’s song, which was statistically significant by 60 dph. This developmental time point is an important one in many respects. First, it represents the beginning of “plastic song”, a phase in which the juvenile’s song rate increases dramatically (Johnson et al., 2002; Zann, 1996) and syllable structure begins to resemble that of the tutor (Kollmorgen et al., 2020). Our results suggest that during this period, pupils spend less time seeking out the song they will eventually sing, perhaps because their efforts have shifted toward practicing rather than listening. The shift in preference, toward neighbor’s song, is interesting because it occurred at a time when juvenile males typically transition from spending time with the family unit to seeking contact with other, unrelated birds (Adkins-Regan & Leung, 2006). This transition may be reflected in their song preferences; Fujii et al. (2021) showed that preferences for the father’s song over that of an unfamiliar male began to wane after 60 dph in males but not in females.

The shift in preference at 55 dph may provide information about underlying neural mechanisms. In a previous study (Davis et al., 2019), we mapped and quantified the distribution of oxytocin receptor (OTR) expression during the entire period of vocal development in zebra finches. We found a striking reduction in OTR mRNA at 55 dph in each brain region we looked at: the lateral septum, the auditory forebrain, and two regions containing song control nuclei. This result, together with our current finding of a dramatic shift in song preferences at exactly the same time point, provides correlational evidence that early attraction to father song may depend on OTR expression in some or all of these regions. Given the well-known role of OTR in social attachment and sociality (Ross & Young, 2009), which has been shown even in zebra finches (Goodson et al., 2009; Pedersen & Tomaszycki, 2012), it is possible that this receptor contributes to socially-mediated vocal learning by mediating the establishment of early song preferences (Maney & Rodriguez-Saltos, 2016; Theofanopoulou et al., 2017).

### Potential genetic contributions

Because we did not cross-foster the pupils in our study to unrelated parents, we do not know whether the preferences exhibited during the key-pressing assay could be explained by a genetic component. A rich literature dating back several decades has shown clearly that tutor choice depends critically on early social experience (Clayton, 1987; Mann et al., 1991; Mann & Slater, 1995; Williams, 1990; Zann, 1996). Young zebra finches choose to sing the song of a foster father even if the biological father can be heard singing in the room (Immelmann, 1969; Roper & Zann, 2006). Nonetheless, there is recent evidence that certain components of song may have a genetic basis (Lansverk et al., 2019; Mets & Brainard, 2018). Future studies of song preferences should include cross-fostering as part of the design.

### An operant conditioning assay to measure preference while balancing exposure

As part of this project, we developed an operant assay that can be used to measure preference for a stimulus while controlling for exposure to that stimulus. This assay may be useful to others, not only those studying song learning but also any process in which the outcome measure may be confounded by exposure effects. We note that in the present study, we may not have needed to control for exposure effects on learning. Two birds, which were excluded from our main analysis, were reared with the father and then given a choice between the songs of two neighbors in the operant assay. Both ultimately sang father’s song instead of either neighbor’s song. Although there were only two birds in this condition, this result adds an interesting layer to our knowledge about what can and cannot be accomplished by operant tutoring. We know from previous work that if the father is removed at 35 dph and a new, live tutor is provided, pupils will learn primarily the new tutor’s song (Eales, 1985; c.f. Gobes et al., 2019). Although many studies have shown that operant tutoring is effective in birds that are relatively naïve to song (Derégnaucourt et al., 2013; Tchernichovski et al., 1999), our data show that if pupils are able to interact with their father until 35 dph, although they will key-press to hear other songs, they ultimately reject them, choosing instead to sing father’s song. We see many possibilities for testing a variety of hypotheses with this assay and hope that others can use it in their own studies of preference.

## MATERIALS AND METHODS

### Ethics statement

All of our procedures involving handling and experimentation with animals were approved by the Institutional Animal Care and Use Committee at Emory University.

### Finch husbandry

Adult zebra finches were randomly paired to produce offspring for our experiment. These breeding pairs were housed in 14×15×17 inch cages in the animal facility at Emory University. The birds were provided with food and water *ad libitum*. To encourage breeding, we sprayed water inside the cage daily, provided hard-boiled eggs and carrots weekly, and nesting material consisting of timothy grass and burlap as needed. All items in the cage, such as food trays, water baths, and bottles, were arranged symmetrically to discourage offspring from developing a preference for either side of the cage. Within a single room, four breeding pairs were housed together, each pair in its own cage. Birds in any cage could hear the ones in the other cages but could not see them due to white plastic dividers placed between the cages.

### Operant chamber

We used only males for this experiment because in this species, only males sing. Juvenile males (n = 13) were separated from their parents at 37 (IQR: 36, 37) days post-hatch (dph) (Fig. 1A). By this age, juvenile zebra finches can feed themselves (Zann, 1996). Each male was isolated from other birds in a 14×15×17 inch cage placed inside a sound-attenuating booth. In this cage, food and water were provided *ad libitum*. Food, water and other items in the cage were arranged symmetrically to reduce side bias. To provide enrichment, a mirror was centered on the rear wall of the cage.

The cage was equipped with two keys (Fig. 1B, C), placed on opposite walls. Upon being pressed, each key elicited playback of either the father’s song (hereafter “father’s song”) or the song of a neighbor (hereafter “neighbor’s song”) from the room where the juvenile was reared. One of the keys had a higher likelihood of eliciting playback of father’s song while the other key had a higher likelihood of eliciting playback of neighbor’s song. Whether the left or right key was associated with father’s or neighbor’s song was balanced across subjects. The keys were connected to a computer via a National Instruments Board USB-6501 (National Instruments, Austin, TX, USA) or an Arduino UNO board (Arduino LLC, Somerville, MA, USA). The computer ran the software SingSparrow!, which we wrote, to control the responses of the keys and to log the presses on them (code available in Supplemental Materials). Each cage had two speakers, each one paired with one key. We used speakers of two models (LS-300, AudioSource, Portland, OR, USA; Logitech Z200, Newark, NJ, USA); in each cage, the two speakers were of the same model.

Operant conditioning proceeded daily until the bird reached 89 dph (IQR: 87, 90) (Fig. 1A), when song is crystallizing, or taking its final form (Zann, 1996). Thus, operant conditioning proceeded throughout the majority of the period when the zebra finches practiced singing.

### Neighbor’s song

In order to control for any effects of exposure on vocal learning, we selected the song of a neighbor that the juvenile heard about the same number of times as he heard his father’s song while in the breeding room. To identify this neighbor, we estimated the singing rates of each of the four adult males in the room during the time that the juvenile was housed there—from hatching to 35-40 dph. We obtained 20-45 recordings of sounds in the room on random days and times of the day. Each recording lasted 10 minutes. Because male zebra finches sing only one song type, which is usually distinct from the song types of other males, we were able to detect each event of singing by each male just by listening to the recordings or looking at their spectrograms. We defined an event of singing as a continuous bout of song lasting for 3 seconds or less. We chose this threshold because it corresponded to the duration of the shortest bout of singing in a sample of the recordings. The average number of events per recording was our estimate of singing rate. To estimate the uncertainty in our estimate for each male, we generated an empirical sampling distribution of singing rates by bootstrapping the number of events per recording (number of bootstraps = 10,000). We took this uncertainty into account when selecting the neighbor with the song that was used in our experiment; among the three neighbors, we selected the bird for which the sampling distribution overlapped the most with that of the father.

### Playback stimuli

For each pupil, the playbacks consisted of one recording of father’s song or one recording of neighbor’s song. Each recording comprised two consecutive song motifs, or units of song, from the song of the corresponding bird. Motifs consist of a series of distinct sounds, known as syllables, that are always sung in the same order by a bird (Zann, 1996). To record the motifs, we separated the male from its female partner for 20 minutes. We then reunited the pair, which prompted the male to sing to the female. The song was recorded using a TASCAM DR-7MKII recorder. The stimulus was high-pass filtered at 800 Hz to eliminate low-frequency noise produced by the recording system. When Logitech Z200 speakers were used (see Operant Chamber), we applied an equalization preset to the stimuli to control for frequency-dependent distortions caused by these speakers.

### Reinforcement schedule

To test whether preference predicts learning, we needed to control the amount of exposure to each song. Otherwise, a correlation between learning and preference may result from increased exposure to the preferred song. As described above, the juveniles’ exposure to father’s and neighbor’s song was roughly equal while the juvenile lived in the breeding room. In addition, we controlled exposure during the operant conditioning phase of the study. To do so, we designed a reinforcement schedule that allowed us to detect a preference for either song, balance exposure to each song, and limit exposure to 30 playbacks of each song per day. This quota of playbacks was chosen to prevent detrimental effects of overexposure on learning (Tchernichovski et al., 1999). Each key in the operant conditioning cage could play both songs, but each had a higher probability of playing either the father’s or neighbor’s song. The probabilistic schedule allowed the birds to play both songs throughout the session while still indicating their preference for one of the songs. Once the quota of their preferred song was reached, the pupils could play only the other song by pressing either key, until its quota was also reached. Once the quotas of both songs were reached, pupils could not elicit any more playbacks that day. The keys were reset the following morning at lights-on.

The probability that a key would play its associated song changed throughout each daily session to prevent the preferred song from being played many more times than the other (Figure 2). At the beginning of the session, if the bird pressed only the key associated with its preferred song, the probability of playing that song was 0.75. In other words, out of every four presses, three resulted in playback of the preferred song and one in playback of the other song (Figure 2). The probability was this high at the beginning of each day to strengthen the association between the key and the song. After 12 consecutive presses, the probability went down to 0.67, and to 0.5 after 12 more presses (Figure 2).

When the probability of playing the preferred song was 0.5, the association between the keys and the songs was maintained because under such a scenario the other key was programmed to never play the preferred song (Figure 2). That key played only the song with which it was associated, until that song had been heard that day the same number of times as the preferred song. In this way, the probability of hearing the associated song for each key was always much higher than hearing the other song.

When a pupil switched keys, the first press after the switch always resulted in playback of the song associated with the newly pressed key, regardless of how many times that song had been played. This rule was introduced to help the bird learn the associations between the keys and the songs. Moreover, if the bird did not have a preference and constantly switched between keys, exposure was naturally balanced by playing the song associated with the key being pressed. If the bird had a preference, then the 30 playbacks of the preferred song were exhausted first. Because of the probabilistic contingencies, by the time the preferred song was exhausted, the other song had been played at least 19 times, and therefore it did not take many presses to end the session. After the end of the session, the keys became silent.

### Reconstructing developmental trajectories of song preference

Logs of key presses were cleaned in four steps before they were used to estimate trajectories of song preference. First, we deleted days in which the pupils had not exhausted the quota of both songs. These days occurred mostly at the beginning of the experiment, while the birds were habituating to the cage and learning the operant task. Second, we removed presses that occurred after the quota of the preferred song for that day was exhausted, because at that point the pupils no longer had a choice between the two songs. We also removed presses that occurred within three seconds of a previous press because these presses were unlikely to be independent of the first. We had programmed the keys not to play more than one song within three seconds of a press, so these extra presses did not elicit playback.

To measure preference for either song, we calculated the daily proportion of presses on the key associated with that song. For pupils with strong preferences, we needed to rule out side biases. We did this by applying a reversal, in other words, reversing the associations between the keys and the songs. Reversals were applied only once per pupil and occurred during the night, in between sessions of operant conditioning. We considered that presses made after a reversal do not indicate preference until the pupils have learned the new contingencies of the keys. We predicted that during learning, the pupil would gradually switch from key to the other. As a result, the proportion of presses for the key associated with the preferred song would immediately drop after the reversal, but gradually increase over the next days. We removed data generated immediately after the reversal and for the duration of the increase (Fig. S1). The duration was determined upon visual inspection of the scatterplot of proportion of presses versus days after reversal. Data were not removed if we did not see a change in proportion of presses for father’s song, which we assumed was explained by learning of the new contingencies happening within a day. We planned to remove all data collected from pupils that did not switch keys after the reversal, which would have indicated side bias, but all birds that underwent a reversal did switch.

To reconstruct the average trajectory of preference for father’s song, we fitted a generalized additive model to our data (Wood, 2011). The dependent variable was preference for the father’s song and the independent variable was age. Bird identity was modelled as a random effect. To constrain predicted values to the interval [0,1], we modelled the dependent variable using a binomial distribution. The relationship between preference and age was modelled using a thin-plate regression spline (Wood, 2003). The model was fitted using the library *mgcv* (Wood, 2017) in R (R Core Team, 2019). *Mgcv* produced a 95% confidence interval of the trajectory by multiplying the standard error of the trajectory by two, subtracting this result from the trajectory to find the lower bound, and summing it to the trajectory for upper bound (Wood, 2017). At any given age the pupils significantly preferred the father’s song over the neighbor’s song when the trajectory was above 0.5 and the confidence interval excluded that value.

In order to find the maximum preference of each pupil for the song of the tutor it chose (determined by inspecting spectrograms, see below), we first fitted individual trajectories of preference for that song. For each bird, we fitted the trajectories using locally estimated scatterplot smoothing (LOESS) (Cleveland et al., 1992) in R (R Core Team, 2019) (Fig. S1). The degree of smoothing was controlled via the span parameter. For each bird, we recorded the global maximum point of the trajectory and the age associated with it.

### Recording of vocalizations and selection of pupil’s songs

Recordings of the vocalizations of pupils were made between 80–90 dph. By 80 dph, song crystallization is well underway and the song is a reliable proxy for adult song (Johnson et al., 2002; Tchernichovski et al., 2001; Zann, 1996). We used the software Sound Analysis Pro (SAP) (Tchernichovski & Mitra, 2004; Tchernichovski, 2011) to record the vocalizations. This software was developed to automatically record zebra finch vocalizations, but it does not distinguish songs from other vocalizations such as calls. Thus, these recordings included diverse types of vocalizations.

To extract pupils’ songs from the recordings, spectrograms of the recordings were generated in Audacity v 2.2.2 (Audacity Team, 2018) using Hanning windows. The window size used for spectral analysis was 512 samples, with overlap between windows of 50%. The value that we chose for the window size allowed us to clearly distinguish frequency contours in the spectrograms. In addition, the corresponding duration of the spectral window (11.61 ms, at a sampling rate of 44100 kHz) was close, within 2-3 milliseconds, to that used in previous studies of song in zebra finches (Mandelblat-Cerf & Fee, 2014; Moore & Woolley, 2019; Tchernichovski et al., 2000). To account for variation across renditions of song, we used the function “sample” from R (R Core Team, 2019) to randomly select 30 songs per pupil, which is a number that is in line with previous studies of song imitation in zebra finches (Feher et al., 2009; Moore & Woolley, 2019). Each of these songs was retrieved from a random position in a randomly selected recording, using the random sampler in R (R Core Team, 2019). The songs were then retrieved by a human observer who scanned the spectrograms of the recordings, moving forward from the randomly chosen position, to the first song after that position. To train the observer to identify songs, we presented them with exemplars of song and other vocalizations, such as calls, recorded from adult finches in our breeding colony. The observer classified a vocalization as song if: 1) it consisted of a train of sounds each less than one second in duration, separated by intervals of silence less than 250 ms in duration, and 2) consecutive sounds in the train differed in shape, as seen in the spectrogram (Tchernichovski et al., 2000; Zann, 1996). Whenever a recording contained repeats of song, which happened often because zebra finches tend to sing in bouts, the observer selected only one repeat.

### Analysis of acoustic similarity

We used Sound Analysis Pro 2011 (SAP2011) to evaluate acoustic similarity between the songs of tutors and pupils (Tchernichovski et al., 2000; Tchernichovski, 2011). To avoid including silent intervals between syllables in the analysis, we used amplitude thresholds to filter out those intervals. To further eliminate noise, we used a bandpass filter of 800-8000 Hz. In our recordings, this bandwidth included most of the energy in song while excluding prominent bands of noise caused by electronic interference from building appliances. Other settings in SAP 2011 were set at default values, which are calibrated for analysis of zebra finch song (Tchernichovski, 2011).

### Testing for a correlation between preference and imitation

We tested whether maximum preference for father’s song was correlated with the pupil’s average similarity to their father’s song. The correlation was tested using a generative additive model with the package *mgcv* (Wood, 2017) in R (R Core Team, 2019). The model included a beta regression link and controlled for the random effect of brood.

## Supporting information

Code for operant assay

## Acknowledgments

We are grateful to Jocelyne Bachevalier, Rob Hampton, and Phil Wolff for comments on a previous version of the manuscript, and to Mohammad Alhamdan, Konya Badsa, Emily Brown, Matt Davis, Isabel Fraccaroli, Evan Goode, Erik Iverson, Timothy Libecap, Camden MacDowell, Teresa Pan, and Gulrukh Shaheen for technical assistance. We also thank Ofer Tchernichovski for technical advice and Erich Jarvis and Laura Carruth for providing founders for our zebra finch colony. This work was supported by National Institutes of Health 1R21MH105811-01A1 to DLM, National Institutes of Health P50MH100029 to GR, and by the Silvio O. Conte Center for Oxytocin and Social Cognition, National Institutes of Health 2P50MH100023. CAR-S was supported in part by the Howard Hughes Medical Institute through an International Student Research Fellowship. The authors declare no conflicts of interest.

