## Supplemental Figures for "Song preferences predict the quality of vocal learning in zebra finches"

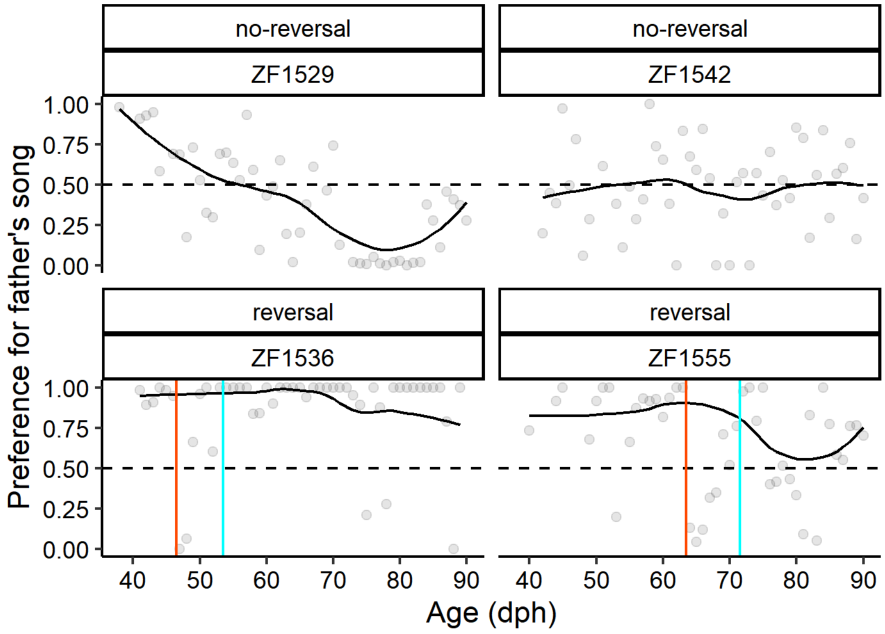


**Fig. S1. Examples of trajectories of preference.** The developmental trajectories of preference are shown for four juveniles in this study. The preference for father's song was calculated as the proportion of presses for the key associated with that song. The gray dots in each plot are the daily preference scores for each bird. A smooth trajectory was calculated by applying LOESS on the datapoints. In birds that showed a strong preference for one of the songs (ZF 1536 and ZF 1555), we applied a reversal (red line) to rule out side biases. The blue line indicates the point at which a human observer considered the bird to have finished switching keys after the reversal. Data between these two vertical lines were not used in LOESS. Despite the original data consisting of proportions, and thus bound to the interval [0-1], LOESS may fit values slightly outside that interval.

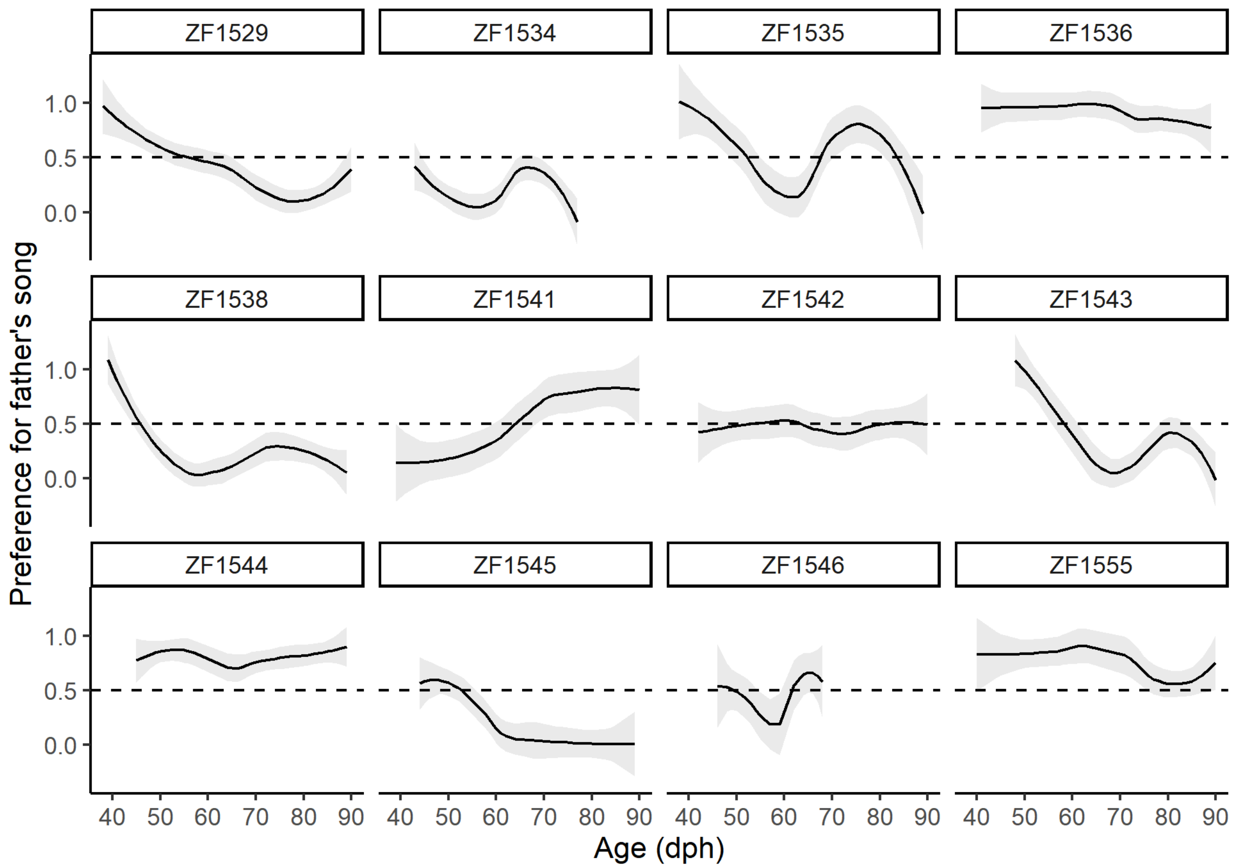


**Figure S2. Individual trajectories of preference for father's song.** The dashed line indicates a preference score of 0.5, at which a bird did not prefer father's or neighbor's song. Scores above 0.5 indicate a preference for father's over neighbor's song. For most birds, preference for father's song peaked early in development and then decreased, shifting to neighbor's song in most birds. The solid lines are smooth trajectories fitted to preferences each day (see Fig. S1) using LOESS. The shaded area is the 95% confidence interval of the trajectory.

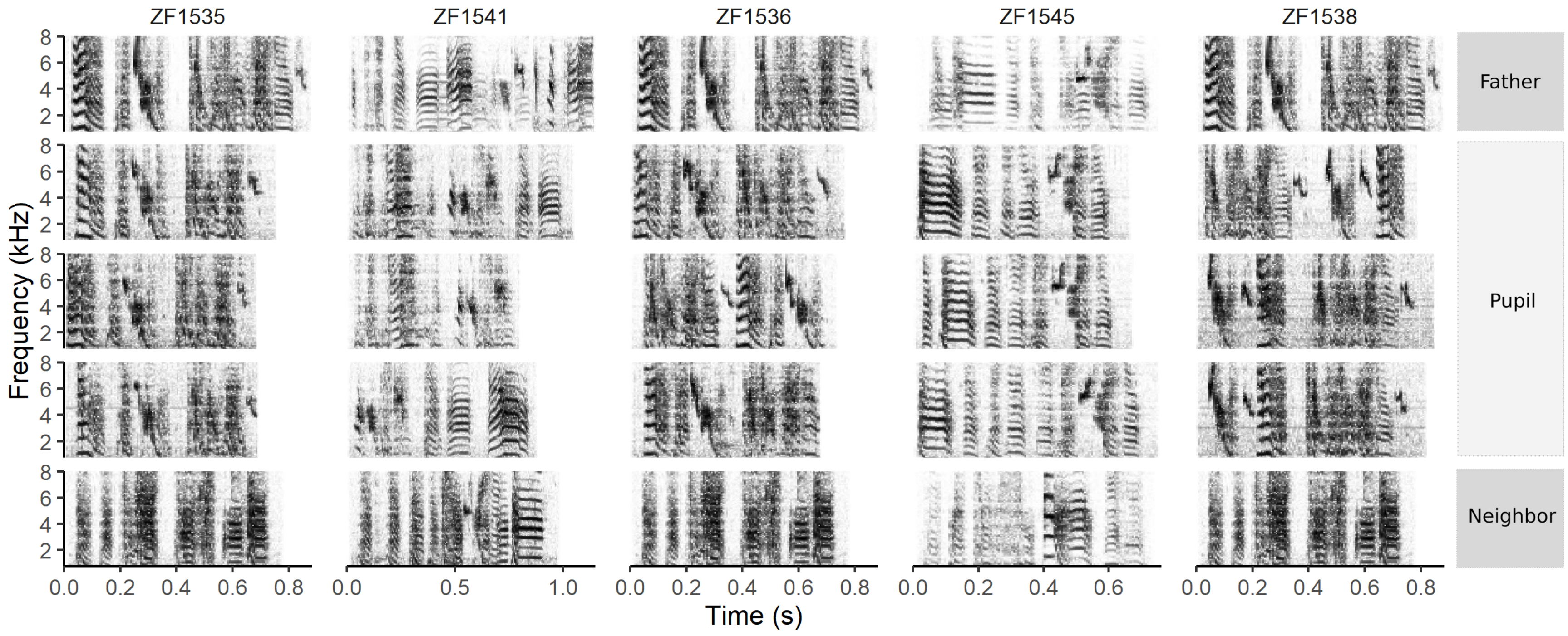


**Fig S3 (part 1). Comparison of spectrograms of father's, pupil's, and neighbor's songs.** In all cases, the songs of the pupils resembled father's song rather than neighbor's song. Some songs were trimmed to eliminate silence and introductory notes. Pupils are organized in decreasing order of average similarity of their songs to their father's song. Five pupils are shown in this panel and five on the next page.

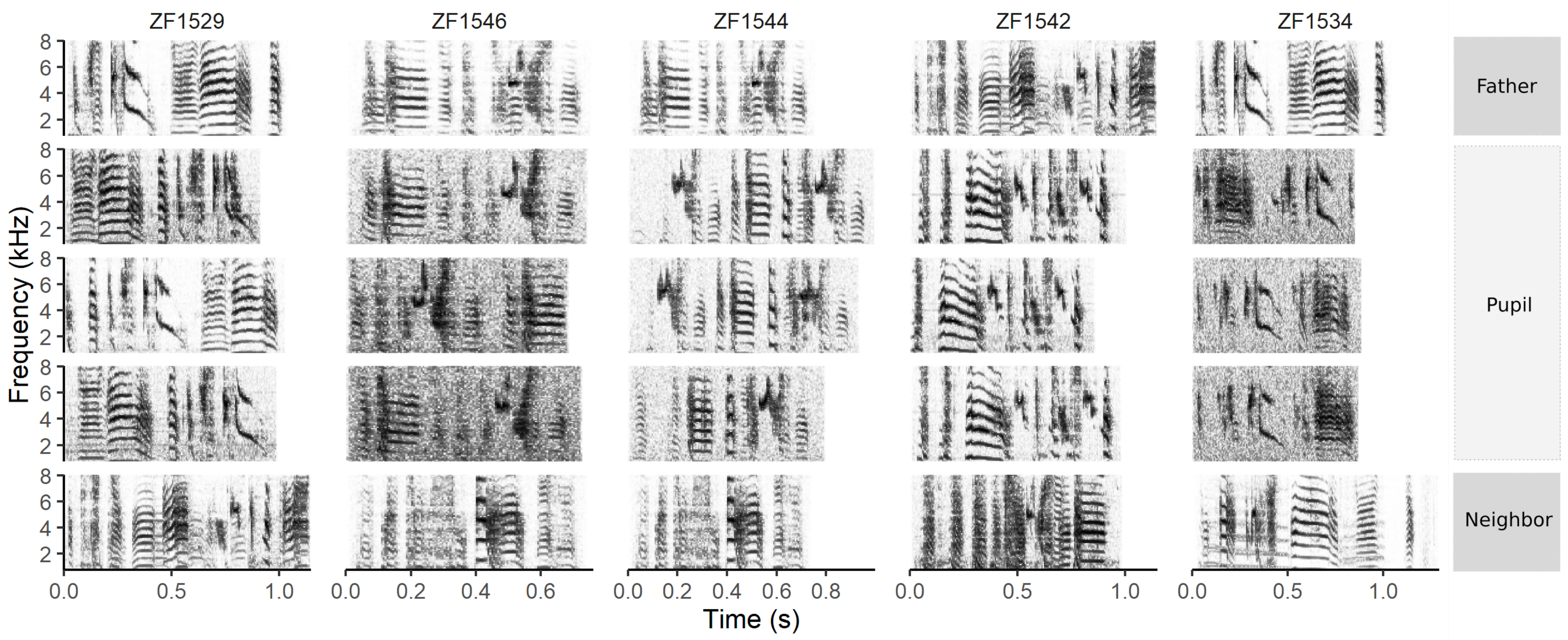


**Fig. S3 (continued from previous page).**

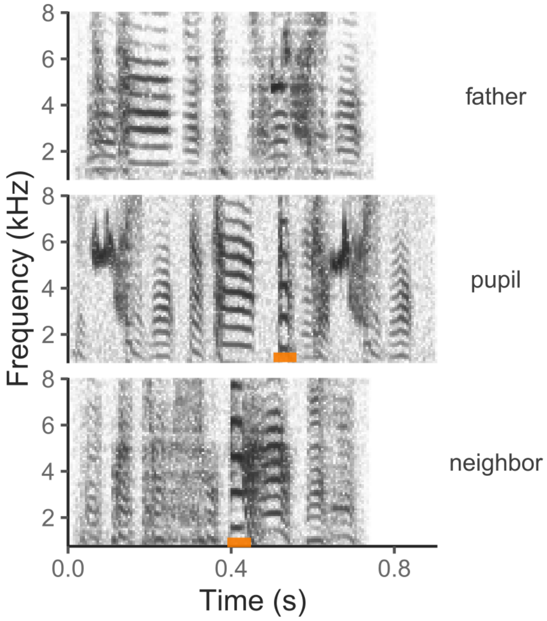


**Fig S4. A syllable copied from neighbor's song.** In the song of one pupil (ZF1544), we found one syllable from the song of the corresponding neighbor. The rest of the song, however, resembled father's song. Orange rectangles mark the syllable in pupil's song and its putative source in the pupil's song.
