## Supplementary material for "Song preferences predict the quality of vocal learning in zebra finches": Code for operant assay: README.docx

**SingSparrow!**

Code written by Carlos Antonio Rodríguez-Saltos, 2014-2015.

Newer versions, written by Evan Goode and Catharine Harris, can be downloaded from "https://github.com/evan-goode/singsparrow-ii".

This program that controls the behavior of playback keys in operant conditioning tests. It balances exposure to both sounds played by the keys while still allowing detection of a preference for a particular key. The keys consist of any type of switch that are connected to the computer through a National Instruments Data Acquisition Device.

The 'Stats' and 'Data Acquisition Toolbox' MATLAB toolboxes are required.

Folder contents

SingSparrow.m – Program written in MATLAB.

parameters.txt – An example file showing how to set the configuration parameters for the program.
