## Supplementary material for "Song preferences predict the quality of vocal learning in zebra finches": Code for operant assay: SingSparrow.m

```
%% SingSparrow!
% Program that controls the behavior of playback keys in operant conditioning tests.
% The program balances exposure to both sounds played by the keys while
% still allowing to detect a preference for a particular key. The keys need to be connected to a National Instruments Data Acquisition Device.

% A newer version of the program can be downloaded from "https://github.com/evan-goode/singsparrow-ii". 
%
% Writen by Carlos Antonio Rodriguez-Saltos, 2014-2015.

% The program  requires the 'Stats' and 'Data Acquisition Toolbox' toolboxes.

%% Import parameters
% The following parameters must be specified by the user in a 'parameter.txt' textfile, which should be placed in the same folder as SingSparrow.m.
%
% [Identifiers] - Parameters that permit to identify the trial.
% bird - Code name of the bird
% booth - Letter desginating the sound isolation booth in which the experiment is carried. Letters could be one from "A", "B", "C", "D", and "E".
% dayTrial - Day number of experiments with the same bird in the same booth
% probe - Whether to run in probe mode. It will output the time elapsed since the program started.
%
% [Key A/B] - Parameters that specify the audio file that will mostly be played by pressing Key A/B.
% folder - Folder containing the audio file. Inside of it, the file should be named "1.wav". If multiple files of the same individual are used, then they should all have unique numbers. The files will be rotated to prevent habituation (see Day schedule below). In our study, we only used one file for individual.
% sound_type - Type of sound in the folder.
% gap - Latency, in seconds, for allowing a key to play sound after the key was pressed.
%
% [Day schedule] - Parameters that specify the schedule of
%                playbacks. 
% Hour start - Hour of the day (24 hour format) at which program starts. Must be an integer.
% Hour end - Hour of the day (24 hour format) at which program ends. Must be an integer.
% Sound change - If multiple playback files are used for the same bird, specifies the interval of time, in minutes, to wait until the sound file is rotated.
% Begin file - Number at which to begin the file rotation.

% [Reinforcement schedule]
% simple - Should the program run in simple mode (no probabilistic schedule)? 0 - No, 1 - Yes.
% MaxPresentations - Maximum number of sound presentations.
% Switch1/2 - Number of playbacks, from onset of experiment, at which the program changes the probabilities of playing song A or B by pressing either key.
% Prob1-3 - Probabilities of playing associated song at each stage in the reinforcement schedule.

% [DAQ] - Specifies the name of the DAQ device and which channels will be
% in use for Key A and Key B. The name of the device can be seen by opening
% the program NI-DAQmx while the DAQ card is connected to the computer. The
% name is usually 'Dev1' or 'Dev2'. The number that identifies each input
% line are labeled in the DAQ itself. For example, the line labeled as
% 'P1.7' corresponds to the line 7 of the port 1. In the parameter file,
% both the port and the line numbers have to be specified as in the
% following example: LineA = 'Port1/Line3'.
% device - Name of the device.
% LineA/B - Input lines of the device assigned to Key A or B, respectively.

% [Sound recorder] - Parameters for setting a sound recording, which will record any sound in the operant booth (microphone needed) when a key is pressed.
% recording - Should recorder be activated? 0 - No, 1 - Yes.
% rate - Sampling rate in Hz.
% bit - Bit depth.
% Buffer - Number of seconds to record before key is pressed.
% After trigger buffer - Number of seconds to record after key is pressed.

% Import datafile 'parameters.txt'
clear all

formatSpec = '%*s bird = %s booth = %s dayTrial = %f probe = %f %*s %*s folder = %s sound_type = %s gap = %f %*s %*s folder = %s sound_type = %s gap = %f %*s %*s Hour start = %f Hour end = %f Sound change = %f Begin file = %f %*s %*s simple = %f MaxPresentations = %f Switch1 = %f Switch2 = %f Prob1 = %f Prob2 = %f Prob3 = %f %*s %*s device = %s LineA = %s LineB = %s %*s %*s recording = %f rate = %f bit = %f Buffer = %f After trigger buffer = %f';
a = fopen('parameters.txt', 'r');
datafile = textscan(a, formatSpec, 'Delimiter', '\n', 'CollectOutput', true);

fclose(a);
clear a formatSpec;

% Load variables from imported datafile
bird = char(datafile{1}(1));
booth = char(datafile{1}(2));
dayTrial = double(datafile{2}(1));
setProbe = logical(datafile{2}(2));

soundA = char(datafile{3}(1));
filesA = dir(soundA);
filesA = filesA(3:end); % eliminates files '.' and '..' from array
numberFiles = length(filesA); % counts the number of files in folder
sounds(1).name = char(datafile{3}(2));
gap(1).limit = double(datafile{4}(1));

soundB = char(datafile{5}(1));
filesB = dir(soundB);
filesB = filesB(3:end); % eliminates files '.' and '..' from array
sounds(2).name = char(datafile{5}(2));
gap(2).limit = double(datafile{6}(1));

startRun = double(datafile{6}(2));
minStart = startRun * 60;
endRun = double(datafile{6}(3));
soundChange = double(datafile{6}(4));
beginFile = double(datafile{6}(5));
currentFile = beginFile - 1; % Making the current file be a different number than beginFile engages the program to load the first file because the program 'thinks' that an update is needed (section Import Files).

simple = logical(datafile{6}(6));
if ~simple % Load the following variables only if simple schedule is deactivated.
    MaxPresentations = double(datafile{6}(7));
    Switch1 = double(datafile{6}(8));
    Switch2 = double(datafile{6}(9));
    Prob1 = double(datafile{6}(10));
    Prob2 = double(datafile{6}(11));
    Prob3 = double(datafile{6}(12));
    Compensation = 0;
end

device = char(datafile{7}(1));
LineA = char(datafile{7}(2));
LineB = char(datafile{7}(3));

setRecorder = logical(datafile{8}(1));
if setRecorder
    recorder.rate = double(datafile{8}(2));
    recorder.bit = double(datafile{8}(3));
    recorder.recTimeBuf = double(datafile{8}(4));
    recorder.recTimeTrigg = double(datafile{8}(5));
end

clear datafile

%% Set counters
nKeys = 2; %Specifies the number of keys, which almost always will be 2. However, having this variable will be useful when the program is upgraded to do more complex experiments.
lastKeySong = 0; %Tells which key was pressed last. 0- No key was pressed, 1- Key A, 2- Key B.

% Generate counters that register the number of times that each key is pressed.
for m = 1:nKeys
    keys(m).pos = 0; % This counter registers the number of times that each key has been pressed and the program sent the command to play song.
    keys(m).total = 0; % This counter registers the number of times each key is pressed without necesarilly having played song
end

% Generate a counter that registers the number of times that any key has been pressed
output.Press = 0;

% Generate counters that register how much time has passed since a key was
% pressed. The sound will not be played when a key is pressed again within
% a certain time of the last time that it was pressed and a command was
% send to play song. That time is defined by the user in the paremeters
% file.
for m = 1:nKeys
    gap(m).state = 0;
    gap(m).tictoc = 0;
end

for m = 1:nKeys
    sounds(m).counter = 0;
end
%% Setting the probe
% Probe is a variable that grows with each iteration of the program. It is useful to know when did an error, if any, occur.
if setProbe
    probe = 0;
end
%% Set the schedule
% This portion of the code sets the schedule that will be used by the keys
% to play the sounds. It balances exposure to both sounds while still setting a difference in the
% probability with which each key plays any of the sounds.

% A structure with 2 field, one for each key, and 4 single-row arrays is built. Each array
% represents a stage during the trial. The number of cells in each array
% depends on the values of Switch1, Switch2, and MaxPresentations, where
% such number is equal to Switch1 in the first array, $$(Switch2 - Switch1)$$ in
% the second, and (MaxPresentations - Switch2) in the third.

% The arrays are filled with zeros and twos, where the ones represent the number of times
% that soundA will be played while the twos, the number of times that sound
% B will be played. Such numbers change with each stage of the trial and
% depend on the odds with which each key plays a given sound at any
% stage. In stage 1, the odds of playing SoundA:SoundB upon pressing keyA
% are 3:1. Such odds at each stage will always be reversed for keyB; in
% stage 1, for example, the odds of playing SoundA:SoundB upon pressing
% keyB are 1:3. In stage 2, the same odds will be 2:1 upon pressing keyA,
% and in stage 3 they will be 1:1.

% To fill the arrays, first a series of fixed ones and twos for each
% corresponding array are built. The number of zeros and twos are
% calculated by multiplying the odds of playing soundA:soundB by the total
% number of cells of the array. The series of ones and twos are
% concatenated to form the array. Finally, the numbers are randomly
% scrambled within each array.

% The purpose of the arrays for the program can be compared to that of a
% magnetic tape in playback device. The program has two "virtual tape heads", one for each key, which
% always sits at the beginning of the tape (cell1 of array1). When a key
% is pressed, the corresponding head reads its tape and plays soundA, if
% the tape reads '1', or soundB, if the tape reads'2'. If, and only if,
% consecutive presses are made on a single key, BOTH tape heads advance. If
% the heads reach the end of the array, then they move to the next array
% (the cassette is changed!). If, however, the bird presses a different
% key, no matter where the heads are, they will be transfered back to the
% beginning of the current array. Also, each time a different key is
% pressed, the program plays the sound associated with that key (soundA for
% keyA and soundB for keyB), regardless of the number stored in the first
% cell of the current array for the key that was pressed.

%Set the schedules for each stage of the trial and key.
if ~simple
    CounterSchedule = 1;
    SchedulePos = [0, Switch1, Switch2, MaxPresentations];
    Presents = [Switch1, Switch2 - Switch1, MaxPresentations - Switch2];
    Repeats = round(times(Presents, [Prob1, Prob2, Prob3]));
    for m = 1:length(Presents)
        Schedule(m).A = [ones(1, Repeats(m)), ...
            2 * ones(1, Presents(m) - Repeats(m))];
        Schedule(m).B = [ones(1, Presents(m) - Repeats(m)), ...
            2 * ones(1, Repeats(m))];
    end
    
    %Randomize the order of numbers within each schedule.
    for m = 1:length(Presents)
        SoundFeedback(m).A = randsample(Schedule(m).A, ...
            length(Schedule(m).A), false);
        SoundFeedback(m).B = randsample(Schedule(m).B, ...
            length(Schedule(m).B), false);
    end
    
    % Integrate all arrays into one matrix per key for easier handling.
    % Each array corresponds to a different row of the matrix.
    % Cells corresponding to inexistent cells in each array are filled
    % with zeros. No non-zero values overlap within the same column.
    % Array 2 begins at column number Switch1 + 1, while Array2 begins at
    % column Switch2 + 1.
    SoundFeedbackMatrix(1).vals = [[SoundFeedback(1).A, ...
        zeros(1, MaxPresentations + 1 - length(SoundFeedback(1).A))]; ...
        [zeros(1,Switch1), SoundFeedback(2).A, ...
        zeros(1, MaxPresentations + 1 - ...
        length(SoundFeedback(2).A) - 12)]; ...
        [zeros(1, Switch2), SoundFeedback(3).A, ...
        zeros(1, MaxPresentations + 1 - length(SoundFeedback(3).A) - ...
        Switch2)]]; % The last column consists of only zeros,
    % to mark the end of trials
    
    SoundFeedbackMatrix(2).vals = [[SoundFeedback(1).B, ...
        zeros(1, MaxPresentations + 1 - length(SoundFeedback(1).B))]; ...
        [zeros(1, Switch1), SoundFeedback(2).B, ...
        zeros(1, MaxPresentations + 1 - ...
        length(SoundFeedback(2).B) - 12)]; ...
        [zeros(1, Switch2), SoundFeedback(3).B, ...
        zeros(1, MaxPresentations + 1 - length(SoundFeedback(3).B) - ...
        Switch2)]]; % The last column consists of only zeros,
    % to mark the end of trials
    
    schedule_key1 = table(SoundFeedbackMatrix(1).vals);
    schedule_key2 = table(SoundFeedbackMatrix(2).vals);
    
    % Save playback matrix to a table for later storage
    writetable(schedule_key1);
    writetable(schedule_key2);

    clear SoundFeedback;
end

%% Set output matrix
%This is the matrix that will be filled-up each time the bird presses a
%key and will eventually be writen into a text file once the trial is over.
%
% The columns contain the following data: Press number, Key that was pressed, Sound that was played, Year, Month, Day, Hour, Minute Second

output.Matrix = [zeros(1,4) clock];
OutputFile = table(output.Matrix);
writetable(OutputFile, strcat('OutputFile.', ...
    datestr(clock, 'yyyymmmdd'), '_', booth, '_Id-', bird,'_File1-', ...
    sounds(1).name,'_File2-', sounds(2).name, '_', num2str(dayTrial), ...
    '.txt'));
output.Song = 0; %

log = [zeros(1) clock]; % Writes a log file table showing that a file 
% was succesfully changed automatically, according to the time specified by the user.
logFile = table(log);
writetable(logFile,strcat('log.', datestr(clock, 'yyyymmmdd'), '_', ...
    booth, '_Id-', bird,'_File1-', sounds(1).name,'_File2-', ...
    sounds(2).name, '_', num2str(dayTrial), '.txt'));

%% Set the sound recorder
% the program can record sound each time that a key is pressed. This is
% particularly useful when a researcher wants to compare the vocalization
% of the bird inside the booth with the sound that was played by the birds
% when it pressed the key.

if setRecorder
    % Set sound recorder parameters
    recorder.triggered = false; % Tells whether a key is pressed or not
    
    % Buffer parameters
    recorder.x = 0;
    recorder.y = 0;
    recorder.z = 0;
    recorder.recTimeNow = 0; % Record now? (?)
    recorder.buffer = 1;
    
    % Start recording
    recordMicrophone = audiorecorder(recorder.rate, recorder.bit,1);
    record(recordMicrophone);
end

%% Set device parameters
% This section sets the required parameters for the Digital Acquisition
% Device (National Instruments) to work. The code will work only with
% digital input devices.
d = daq.getDevices %Get device info

% Set session for the device.
s = daq.createSession('ni');
s.DurationInSeconds = 10;
addDigitalChannel(s, device, LineA, 'InputOnly');
addDigitalChannel(s, device, LineB, 'InputOnly');
s.Channels % Show channel info.

a = [1:2;1:2]'; % Matrix that stores up to 1000 events from the device. It acts as a memory buffer, overwriting values, starting with th the first rows, once the end has been reached. Theh matrix is necessary to know whether a "press" event is associated with the onset of the pressing or with a pressing that is withheld. Probably, 1000 rows is too much for the matrix.

%% Run the program during On hours
masterTime = clock;
tic;
while masterTime(4) < endRun % Runs program until time specified by user (endRun). At this stage, the program can be run continuously only within a single day.
    %% Import sound files
    % The sounds are imported as numeric matrixes contained within the
    % structure array 'sounds(1/2)'. The structure sounds(1) contains the sound
    % data for the playback of Key A while the structure sounds(2) contains
    % that for Key B. Within each structure there is information on the type of
    % sound contained in the file (sound().name) and the number of times that
    % the file has been played in the current trial (sounds().counter).
    % The sounds loaded into sounds(1) are always played to the Left output
    % channel, while the sounds loaded into sounds(2) are always played to the
    % Right output channel. Note that the orientation of the computer output
    % channels may not be the same as the physical orientation of the speakers
    % playing the sound inside the booth. For the sake of simplicity, it is
    % recommended to always associate KeyA with the Left ouput channel and KeyB
    % with the Right output channel.
    
    % get directory files for sound A and B
    % currentFile will get
    % updated every 15 minutes
    timeFile = mod(ceil((((masterTime(4) * 60) + masterTime(5)) - minStart) / soundChange) + beginFile - 2, numberFiles) + 1 ; % Tells the program what file whould be played, based on the time of the day.
    
    if ~(currentFile == timeFile) % Update sound
        [sounds(1).data, sounds(1).Fs] = audioread(strcat('./',soundA, '/', filesA(timeFile).name));
        if size(sounds(1).data, 2) == 1 % If sound is mono, this command converts it to stereo
            sounds(1).data = [sounds(1).data sounds(1).data];
        end
        
        [sounds(2).data, sounds(2).Fs] = audioread(strcat('./',soundB, '/', filesB(timeFile).name));
        if size(sounds(2).data, 2) == 1 % If sound is mono, then duplicate the channels
            sounds(2).data = [sounds(2).data sounds(2).data];
        end
        
        currentFile = timeFile;
        
        % Write that change of file was succesful
        log = [log; [currentFile, clock]];
        logFile = table(log);
        writetable(logFile,strcat('log.', datestr(clock, 'yyyymmmdd'), '_', booth, '_Id-', bird,'_File1-', sounds(1).name,'_File2-', sounds(2).name, '_', num2str(dayTrial), '.txt'));
    end
    
    for n = 1:2 % Update cycle of device and sound recorder.
        %% Set clocks and timers
        %  Timer, set by tic/toc functions, have multiple functions in the program. They are used to count how many seconds has
        %  it lapsed from the last key stroke. If consecutive pecks happen too
        %  soon, the program does not play a sound but it records the peck. They
        %  are used to buffer sound when recording, and allow to record sound in
        %  the booth before and after the key press.
        % masterTime logs the time from the clock computer. The time will get
        % updated while the program runs. Setting it here is useful for variables
        % that need to be specified before running the loop.
        if setProbe
            probe = probe + toc
        end
        masterTime = clock;
        %% Update recorder
        if setRecorder
            recorder.recTimeNow = recorder.recTimeNow + toc; % How much time has the recorder been recording
            
            if recorder.recTimeNow >= recorder.recTimeTrigg && recorder.triggered % What to do if enough time (specified by user) has passed since a key was triggered to play a sound.
                recorder.time = datestr(clock, 'yyyymmdd_HHMMSS');
                
                % Reset recorder and write buffers
                stop(recordMicrophone);
                recorder.recTimeNow = 0;
                
                if recorder.buffer == 1
                    sound2write = [recorder.x;recorder.y;recorder.z;getaudiodata(recordMicrophone)];
                elseif recorder.buffer == 2
                    sound2write = [recorder.y;recorder.x;recorder.z;getaudiodata(recordMicrophone)];
                end
                
                audiowrite(strcat(booth, recorder.time, '.wav'),sound2write,recorder.rate);
                recorder.x = 0;
                recorder.y = 0;
                recorder.z = 0;
                recorder.buffer = 1;
                
                clear recordMicrophone;
                recordMicrophone = audiorecorder(recorder.rate, recorder.bit,1);
                record(recordMicrophone);
                
                recorder.triggered = false;
            end
            
            if recorder.recTimeNow >= recorder.recTimeBuf && ~recorder.triggered % Specify what to do when recorder buffer gets full, according to the time specified by user.
                stop(recordMicrophone);
                recorder.recTimeNow = 0;
                
                % The recorder works with two memory buffers. It alternates
                % the loading on each buffer.
                if recorder.buffer == 1
                    recorder.x = getaudiodata(recordMicrophone);
                    recorder.buffer = 2;
                elseif recorder.buffer == 2
                    recorder.y = getaudiodata(recordMicrophone);
                    recorder.buffer = 1;
                end
                
                clear recordMicrophone;
                recordMicrophone = audiorecorder(recorder.rate, recorder.bit,1);
                record(recordMicrophone);
            end
        end
        
        %% Update gap state
        % Enable a key after enough time (in seconds) has passed after its
        % inactivation
        for m = 1:nKeys
            if gap(m).state == 1
                gap(m).tictoc = gap(m).tictoc + toc;
                if gap(m).tictoc >= gap(m).limit
                    gap(m).state = 0;
                end
            end
        end
        tic; % tic has to be here so that on the next cycle of updates of device and sound recorder all variables that depend on toc get accurate updates.
        %% Update device
        a(n,:) = inputSingleScan(s);
        pressed = a(n,:); % Gets the actual value for each input channel
        
        %% Detect and respond to key presses
        if sum(pressed) ~= nKeys % Check whether a key has been pressed
            for j = 1:length(pressed) % Scan the signal sent by each key
                if a(mod(n - 2,2) + 1,j) == 1 && a(n,j) == 0 % Check whether the press is an onset of press.
                    % Count the key press
                    output.time = clock;
                    keys(j).total = keys(j).total + 1; % Counts the number of times that each key was pressed
                    output.Press = output.Press + 1; % Transfers info to output
                    output.Key = j; % Transfers information on which key was pressed
                    
                    % Play the appropiate sound if key was pressed
                    if gap(j).state == 0
                        if ~simple
                            if CounterSchedule < 4
                                if setRecorder
                                    % Start recording sound if key was
                                    % triggered
                                    stop(recordMicrophone);
                                    recorder.recTimeNow = 0;
                                    
                                    if recorder.triggered
                                        recorder.z = [recorder.z;getaudiodata(recordMicrophone)];
                                    else
                                        if recorder.buffer == 1
                                            recorder.x = getaudiodata(recordMicrophone);
                                            recorder.buffer = 2;
                                        elseif recorder.buffer == 2
                                            recorder.y = getaudiodata(recordMicrophone);
                                            recorder.buffer = 1;
                                        end
                                    end
                                    
                                    clear recordMicrophone;
                                    recordMicrophone = audiorecorder(recorder.rate, recorder.bit,1);
                                    record(recordMicrophone);
                                    
                                    recorder.triggered = true; % The key was pressed, thus, it triggers the sound playback and recorder
                                end
                                
                                % Activate gap
                                gap(j).state = 1;
                                gap(j).tictoc = 0;
                                
                                if Compensation == 0 % Refers to the compensation after a sound has been played MaxTrials/2 times.
                                    if lastKeySong ~= j % If the bird has switched from key, the sound played by the new key is that which has a higher probability of being delivered after pressing that key.
                                        sound2Play = j;
                                        for k = 1:nKeys
                                            keys(k).pos = SchedulePos(CounterSchedule);
                                        end
                                    else
                                        keys(j).pos = keys(j).pos + 1;
                                        sound2Play = SoundFeedbackMatrix(j).vals(CounterSchedule,keys(j).pos);
                                    end
                                    
                                    % If the sound associated with the key is the sound that has been played with the lowest frequency overall, then a compensation schedule starts, which will play only the sound associated with the key until both sounds have the same amount of playbacks.
                                    if sound2Play ~= j && sounds(j).counter < sounds(mod(j,2) + 1).counter
                                        sound2Play = j;
                                    end
                                    
                                    queueSound = sounds(sound2Play).data;
                                    queueSound(:,mod(j,2) + 1) = zeros(size(queueSound, 1),1);
                                    sound(queueSound, sounds(sound2Play).Fs); % Plays the sound
                                    disp(strcat('Playing',32, sounds(sound2Play).name,32, 'at',32,datestr(clock, 'yyyy-mm-dd HH:MM:SS'))); % Displays info about the sound just played
                                    
                                    sounds(sound2Play).counter = sounds(sound2Play).counter + 1; % Counts up the number of times a particular song was played
                                    output.Song = sound2Play;
                                    lastKeySong = j;
                                    
                                    if sounds(sound2Play).counter == MaxPresentations / 2 % When a song has been played 30 times, this code disables the playback of that song in both keys, and only enables the playback of the other song.
                                        soundIndexes = 1:2; soundIndexes(sound2Play) = [];
                                        Compensation = soundIndexes; % For this to work, only two sounds can be played in each booth. In conspecific versus heterospecific song preference studies, only two sounds are played in each booth.
                                    end
                                    
                                    % Switches to next part of the schedule, based on number of consecutive presses of a particular key
                                    if SoundFeedbackMatrix(j).vals(CounterSchedule, keys(j).pos + 1) == 0
                                        %if any(Presses.Song == SchedulePos(2:end)) % Switches to next schedule when keys have been pressed a certain number of times (it counts presses for both A and B)
                                        CounterSchedule = CounterSchedule + 1;
                                        for k = 1:nKeys
                                            keys(k).pos = SchedulePos(CounterSchedule);
                                        end
                                    end
                                else
                                    keys(j).pos = keys(j).pos + 1;
                                    sound2Play = Compensation;
                                    
                                    queueSound = sounds(sound2Play).data;
                                    queueSound(:,mod(j,2) + 1) = zeros(size(queueSound, 1),1);
                                    sound(queueSound, sounds(sound2Play).Fs); % Plays the sound
                                    disp(strcat('Playing',32, sounds(sound2Play).name,32, 'at',32,datestr(clock, 'yyyy-mm-dd HH:MM:SS'))); % Displays info about the sound just played
                                    
                                    sounds(sound2Play).counter = sounds(sound2Play).counter + 1; % Counts up the number of times a particular song was played
                                    output.Song = sound2Play;
                                    
                                    if sounds(sound2Play).counter == MaxPresentations / 2 % By this time, both songs would have been played 50 times each.
                                        CounterSchedule = 4;
                                        for k = 1:nKeys
                                            keys(k).pos = SchedulePos(CounterSchedule);
                                        end
                                    end
                                end
                            end
                        else
                            if setRecorder
                                % Start recording sound if key was
                                % triggered
                                stop(recordMicrophone);
                                recorder.recTimeNow = 0;
                                
                                if recorder.triggered
                                    recorder.z = [recorder.z;getaudiodata(recordMicrophone)];
                                else
                                    if recorder.buffer == 1
                                        recorder.x = getaudiodata(recordMicrophone);
                                        recorder.buffer = 2;
                                    elseif recorder.buffer == 2
                                        recorder.y = getaudiodata(recordMicrophone);
                                        recorder.buffer = 1;
                                    end
                                end
                                
                                clear recordMicrophone;
                                recordMicrophone = audiorecorder(recorder.rate, recorder.bit,1);
                                record(recordMicrophone);
                                
                                recorder.triggered = true; % The key was pressed, thus, it triggers the sound playback and recorder
                            end
                            
                            % Activate gap
                            gap(j).state = 1;
                            gap(j).tictoc = 0;
                            
                            % Play sound
                            sound2Play = j; % If the program is run on a simple schedule, then the only sound that will be played upon pressing a key is the asound associated with that key.
                            
                            queueSound = sounds(sound2Play).data;
                            queueSound(:,mod(j,2) + 1) = zeros(size(queueSound, 1),1); % Fill with zeros the channel (left or right) that will not play the sound
                            sound(queueSound, sounds(sound2Play).Fs); % Plays the sound
                            disp(strcat('Playing',32, sounds(sound2Play).name,32, 'at',32,datestr(clock, 'yyyy-mm-dd HH:MM:SS'))); % Displays info about the sound just played
                            
                            sounds(sound2Play).counter = sounds(sound2Play).counter + 1; % Counts up the number of times a particular song was played
                            output.Song = sound2Play;
                            lastKeySong = j;
                        end
                    end
                    %% Write press event
                    output.Matrix = [output.Matrix; [output.Press, output.Key, output.Song, currentFile, output.time]];
                    OutputFile = table(output.Matrix);
                    writetable(OutputFile, strcat('OutputFile.', datestr(clock, 'yyyymmmdd'), '_', booth, '_Id-', bird,'_File1-', sounds(1).name,'_File2-', sounds(2).name, '_', num2str(dayTrial), '.txt'));
                    output.Song = 0;
                end
            end
        end
    end
end

%% End program
log = [log; [0, clock]];
logFile = table(log);
writetable(logFile,strcat('log.', datestr(clock, 'yyyymmmdd'), '_', booth, '_Id-', bird,'_File1-', sounds(1).name,'_File2-', sounds(2).name, '_', num2str(dayTrial), '.txt'));
```
